# Efflux pump regulation and metabolic rewiring define the ceiling of efflux-mediated drug resistance in Mycobacteria

**DOI:** 10.64898/2026.01.04.697598

**Authors:** Shiksha Sharma, Sarika Mehra

## Abstract

Multidrug-resistant efflux pumps play a critical role in antimicrobial resistance, yet the physiological limits of their overexpression remain poorly understood. While gene amplifications and regulatory mutations commonly drive this overexpression, these pumps remain transcriptionally inactive under normal cellular conditions due to their high energetic cost, making it challenging to assess their full impact on resistance evolution and the maximum attainable minimum inhibitory concentration (MIC). To delineate the extent and spectrum of efflux-based resistance, we subjected *Mycobacterium smegmatis mc²155(Msm)* wild-type and efflux pump-deleted strains (Δ*lfrA* and Δ*efpA*) to adaptive laboratory evolution in the presence of a broad-spectrum efflux substrate ethidium bromide (EtBr). Despite prolonged selection, resistance plateaued at 8–16× MIC, revealing an intrinsic ceiling to efflux-based adaptation. Evolved populations integrated multiple layers of adaptation: enhanced efflux kinetics, reduced intracellular EtBr accumulation, and mutations spanning global regulators, ribosomal genes, and membrane components. Notably, deletion of specific pumps (Δ*lfrA)* triggered compensatory upregulation of alternative efflux systems, highlighting functional redundancy. Beyond efflux activation, evolved strains exhibited metabolic reprogramming, including downregulation of the tricarboxylic acid cycle and upregulation of lipid biosynthesis and β-oxidation, which altered NAD⁺/NADH homeostasis and membrane potential. Extending this adaptive evolution approach to *Mycobacterium tuberculosis* H37Ra (*Mtb)* demonstrated that these adaptive constraints and metabolic alterations are conserved across mycobacterial species. Together, these results uncover fundamental limits to efflux evolution and reveal how bacteria integrate regulatory and metabolic plasticity to maximize drug resistance. Targeting these interconnected networks may offer a powerful strategy to contain the rise of multidrug resistance.

**Importance:** Efflux pumps are widely recognized as major contributors to antimicrobial resistance; however, the extent to which resistance can be driven solely by efflux remains unclear. Here, we define the limits of efflux-mediated resistance using adaptive laboratory evolution to ethidium bromide in *Mycobacterium smegmatis* and *Mycobacterium tuberculosis*. We show that resistance reproducibly plateaus despite continued selection and the presence of multiple efflux pumps in the genome, indicating a physiological ceiling to efflux-based adaptation. This limit is achieved through distinct evolutionary trajectories involving regulatory mutations that converge on enhanced efflux activity. Importantly, upregulation of efflux pumps is coupled to extensive cellular remodelling, including metabolic rewiring, and changes in membrane energetics, revealing system-wide trade-offs associated with resistance. Together, our findings demonstrate that efflux-driven resistance is constrained by cellular physiology and shaped by global adaptive costs, thereby refining current models of antimicrobial resistance and identifying vulnerabilities that may be exploited to limit the evolution of resistance.

## Introduction

Antibiotics have revolutionized modern medicine, drastically reducing mortality from bacterial infections and enabling advanced clinical procedures. However, the widespread and often inappropriate use of antibiotics has accelerated the emergence and spread of antimicrobial resistance (AMR), posing a major global health threat. In 2021 alone, AMR was estimated to have caused 4.71 million deaths worldwide, a figure projected to rise to 8.2 million annually by 2050 (1,2). Bacteria develop resistance to antibiotics through various mechanisms, including the alteration of drug targets (3), upregulation of drug-inactivating enzymes (4,5), modification of membrane permeability to reduce drug influx (6), and overexpression of efflux pumps that expel antimicrobial agents from the cell (7). Among these, efflux pumps and their regulatory networks are particularly critical in the development of multidrug resistance (MDR), as they can provide cross-resistance to a broad spectrum of antibiotics (8). Mutations in efflux systems, including both pumps and their regulators, have been documented in clinical isolates of pathogens such as *Mycobacterium tuberculosis* (9–11).

Efflux pumps are energetically costly to operate, and under normal conditions, their expression is tightly regulated (12). However, when bacteria are exposed to antimicrobial agents, certain pumps can be induced or derepressed through specific mutations, enabling the cells to expel toxic compounds more efficiently (13,14). It is important to note that not all efflux systems are inducible; in some cases, overexpression arises from mutations that disrupt transcriptional repression (13). Hence, studies based on mere induction of the efflux pumps in the presence of drugs would not provide a holistic picture of the effect of efflux pumps. While numerous studies have highlighted the role of efflux pumps in contributing to antibiotic resistance (14–19), the upper limit of resistance achievable solely through efflux upregulation remains largely unexplored.

To probe this threshold, adaptive laboratory evolution (ALE) presents a powerful approach. ALE allows for the stepwise selection of resistant populations under defined selective pressures, enabling the identification of key efflux systems and their associated regulatory components. Prior ALE studies have demonstrated that the resistance trajectory of a given bacterial population is influenced by both intrinsic factors (e.g., genome architecture) and extrinsic factors (e.g., nature of the drug used for selection) (20–22). While many studies have identified efflux-related mutations in evolved populations (23–28), most of these use clinically relevant antibiotics that exert selection on their primary cellular targets, thereby obscuring the contributions of efflux-based resistance.

To specifically assess the maximal resistance attainable through efflux activity, we selected ethidium bromide (EtBr) as a selective agent for ALE. EtBr is a DNA-intercalating compound and a known substrate of multiple efflux pumps from diverse efflux pump families (15,29,30). Importantly, the primary resistance mechanism against EtBr involves active efflux rather than target modification, making it a suitable compound for enriching efflux-based resistance phenotypes.

In this study, we employ *Mycobacterium smegmatis* mc2 155 (*Msm*) as a model system to investigate the evolution of efflux-mediated resistance under selective pressure from EtBr. *Msm*, is a widely used surrogate for dissecting mycobacterial physiology and drug resistance mechanisms due to its substantial genomic and physiological similarity with *Mycobacterium tuberculosis* (31). We further extend the evolution framework to *Mycobacterium tuberculosis* H37Ra (*Mtb*) in order to define the upper limits of efflux-driven resistance in a clinically relevant mycobacterium. Our study aimed to address two key questions: first, what genetic and transcriptional changes facilitate high-level efflux-mediated resistance under EtBr pressure; and second, whether efflux alone is sufficient to confer this resistance or if additional cellular adaptations are required. This comprehensive approach aims to uncover not only the key efflux systems but also broader cellular responses that contribute to antimicrobial resistance, providing insights into potential targets for future therapeutic interventions.

## Results

### Evolution in the presence of ethidium bromide leads to efflux-mediated resistance

To examine how exposure to a broad efflux pump substrate shapes the evolution of antimicrobial resistance, we subjected *Msm* to adaptive laboratory evolution under gradually increasing EtBr concentrations. To further dissect the contribution of efflux systems, we included deletion mutants lacking two key efflux pumps—Δ*lfrA* (MSMEG_6225), whose loss causes hypersensitivity to EtBr (32) and, Δ*efpA* (MSMEG_2619), which encodes an efflux pump associated with elevated resistance to multiple antituberculosis agents (11,18). Wild-type and efflux pump-deficient strains were evolved independently under gradually increasing EtBr concentrations from 0.25×MIC over 50 generations (Fig 1A & B). For each background, three replicate lineages (E1-E3) were evolved, yielding nine independently evolved populations in total.

**Fig 1:**
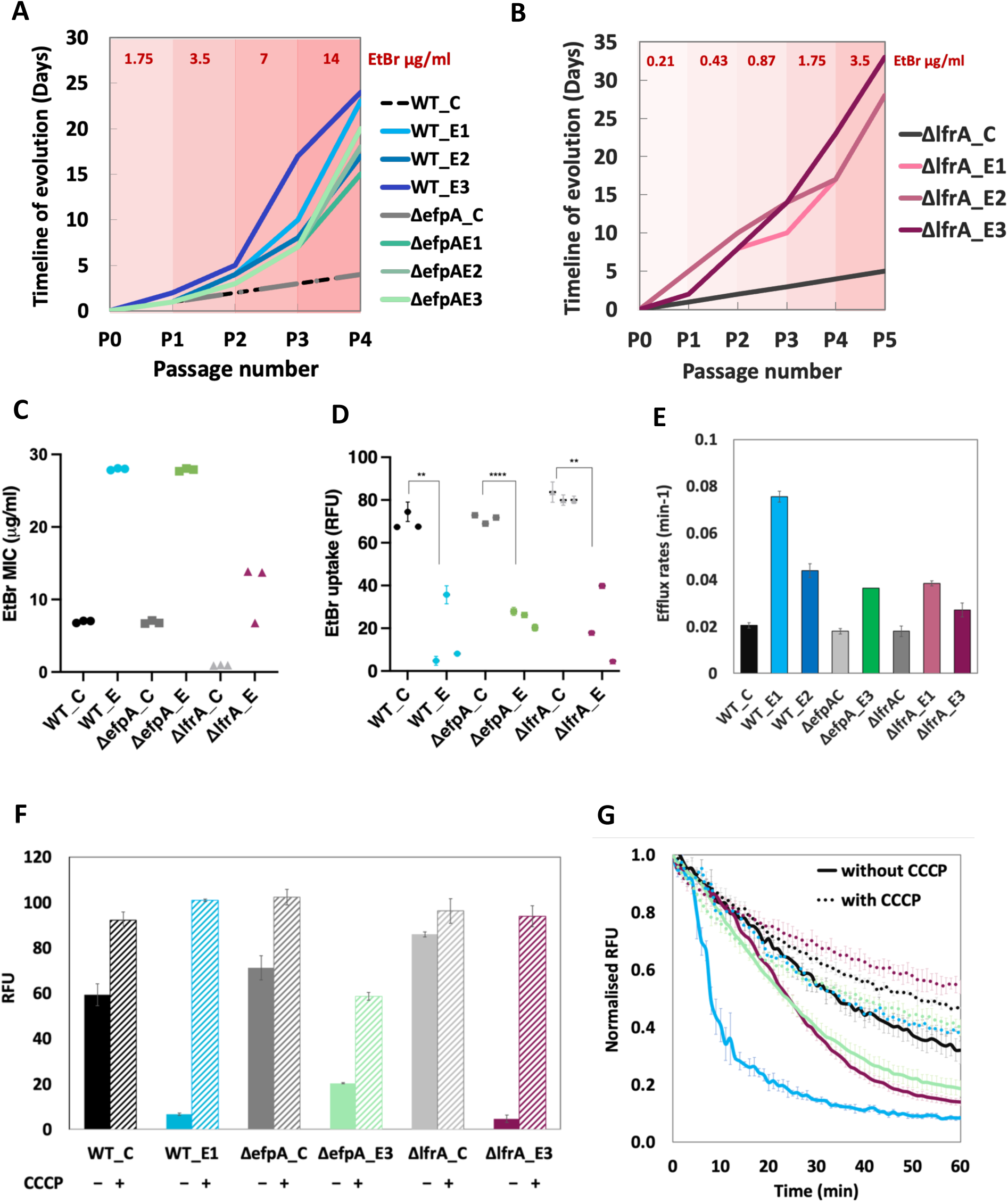
Evolution of Msm strains in the presence of EtBr and their phenotypic characterization. **(A)** Timeline of the evolution experiment for WT and ΔefpA Msm cells. E1, E2, and E3 are respective independent lineages. C represents the control cells evolved without EtBr selection pressure. **(B)** Timeline of the evolution experiment for ΔlfrA cells. ΔlfrA E1, E2, and E3 are three independent lineages. ΔlfrA_C are cells evolved without EtBr pressure. **(C**) MIC of EtBr for the evolved and control strains. WT_E, ΔlfrA_E, and ΔefpA_E represent the MIC values of 3 colonies isolated from three independent evolved lineages (E1-E3) from individual parent strains. WT_C, ΔlfrA_C, and ΔefpA_C represent the corresponding control populations evolved without EtBr. MIC values were confirmed using three to five biological replicates**. (D)** EtBr uptake in control and evolved strains was measured at 3.5μg/ml EtBr over 60 minutes. End-point readings are plotted here. Each dot represents an individual colony from 3 independent evolved lineages, E1, E2, and E3, from wildtype and efflux-deleted parent strains**. (E)** EtBr Efflux rates for evolved strains and their respective controls, calculated using nonlinear regression (one-phase exponential decay) in GraphPad Prism. Values represent the average of three biological replicates. **(F)** The EtBr accumulation in wild-type controls and evolved strains at an EtBr concentration of 3.5 μg/ml for 60 minutes. CCCP at 10 μg/ml was used as a control to inhibit efflux and to restore accumulation. End-point readings are plotted **(G)** EtBr efflux kinetics in wild-type control and evolved strains at 3.5 μg/ml EtBr measured over 60 minutes. CCCP (10 μg/ml) was used as a control to inhibit efflux. Efflux kinetics without CCCP are shown as solid lines and with CCCP are shown as dotted lines. Black represents WT_C, sky-blue is WT_E1, magenta for ΔlfrA_E3 and green for ΔefpA_E. Efflux kinetics for the rest of the strains are shown in Fig S2. Error bars represent the standard deviation calculated from three biological replicates. ** indicates p-value <0.01 and **** indicates p-value <0.0001

The wild-type as well the Δ*efpA* populations grew in the presence of sub-MIC level EtBr levels within 5 days. Beyond this threshold, adaptive trajectories diverged across lineages, with adaptation times varying from 17 to 24 days (Fig 1A-B). Despite these differences, all lineages converged on a similar resistance ceiling of ∼4-8× MIC, suggesting that EtBr resistance evolved independently of genetic background. Increasing the population bottleneck by tenfold (10^7^–10⁸ cells/mL) did not extend this limit (Fig S1), indicating that the adaptive plateau reflects an intrinsic ceiling of efflux-based resistance in *Msm*.

Phenotypic analysis of single colonies isolated from evolved populations revealed a consistent increase in EtBr resistance. Wild-type evolved mutants (WT_E1-E3) exhibited a fourfold increase in MIC (28 μg/ml), relative to the parental strain (Fig 1C). The Δ*efpA*-evolved mutants exhibited a similar fourfold increase, whereas the Δ*lfrA* lineages (Fig 1B) achieved an 8-16 fold higher MIC compared to their parental strain (Fig 1C), highlighting compensatory mechanisms that arise in the absence of *lfrA.* Notably, despite these compensatory adaptations, the absolute MIC of the evolved *ΔlfrA* strains remained lower than that of the evolved wild-type populations.

To probe the physiological basis of resistance, we quantified intracellular EtBr accumulation. Evolved lineages from all genetic backgrounds displayed markedly reduced EtBr fluorescence compared to their respective ancestors (Fig 1D). To test whether this reduction resulted from active efflux rather than altered membrane permeability, we performed EtBr accumulation assays in the presence of Carbonyl Cyanide 3-Chlorophenylhydrazone (CCCP), a known inhibitor of major facilitator superfamily (MFS) efflux pumps in mycobacteria (33). CCCP treatment restored EtBr accumulation to ancestral levels (Fig 1F and S3), confirming that resistance was primarily mediated by energy-dependent efflux activity.

To directly quantify the efflux capacity of the evolved mutants, we saturated the cells with sub-MIC concentrations of EtBr, followed by real-time measurement of efflux kinetics. Among the wild-type evolved strains, WT_E1 exhibited the highest efflux rate as compared to WT_C, as well as to mutants of other genetic backgrounds (Fig 1E and G). Within the Δ*efpA and* Δ*lfrA* evolved lineages, Δ*efpA*_E3 and Δ*lfrA*_E3 displayed higher efflux rates relative to their controls (Fig 1E). The addition of CCCP restored the efflux activity of all the mutants to that of the wild-type (Fig 1G). These findings suggest that while efflux-mediated resistance is a common adaptive strategy, the degree of efflux enhancement varies among lineages, likely reflecting distinct evolutionary trajectories and mutational landscapes. Interestingly, although the Δ*lfrA* evolved mutants exhibit a greater fold increase in EtBr MIC compared to the wild-type evolved strains, their efflux rates were lower. This likely reflects the absence of *lfrA*, a key EtBr efflux pump, limiting export efficiency despite compensatory adaptation. The mechanisms underlying this compensation—whether through upregulation of secondary efflux systems, alterations in membrane permeability, or other compensatory genetic modifications remain to be elucidated. Notably, all evolved strains demonstrated partial inhibition of efflux in the presence of CCCP, consistent with its specific inhibition of proton motive force (PMF)-dependent MFS efflux pumps, while ATP-binding cassette (ABC) transporters likely remain active. Overall, these findings highlight efflux as the dominant resistance mechanism under EtBr selection, with strain-specific variation in the magnitude and mechanistic basis of efflux enhancement.

### Genotypic changes reveal mutations in membrane proteins and transcription factors

To elucidate the genetic basis of resistance in the EtBr evolved lineages, whole-genome sequencing was performed on representative clones from each evolved population. A heatmap representation is shown in Fig 2. In the wild-type lineage WT_E1, a single nucleotide polymorphism (SNP) was identified in the *lfrR* gene (MSMEG_6223), which encodes the transcriptional repressor of the *lfrA* efflux pump. Loss-of-function mutations in *lfrR* are known to lead to constitutive overexpression of *lfrA*, enhancing EtBr efflux (34). WT_E1 also carried a mutation in *MSMEG_1959*, encoding a conserved membrane protein of the uncharacterized UPF0182 family. The WT_E2 strain harboured mutations in both *soxR*, a global transcriptional regulator implicated in oxidative stress responses and efflux regulation, and in *MSMEG_1959*, consistent with the mutation seen in WT_E1. In contrast, WT_E3 displayed a distinct mutation profile, with no overlap in efflux regulator mutations but still retaining a variant in *MSMEG_1959*. Notably, *MSMEG_1959* was the only gene mutated across all three wild-type evolved lineages, suggesting a potential role in adaptation to EtBr exposure, possibly through membrane remodeling or altered permeability.

**Fig 2.**
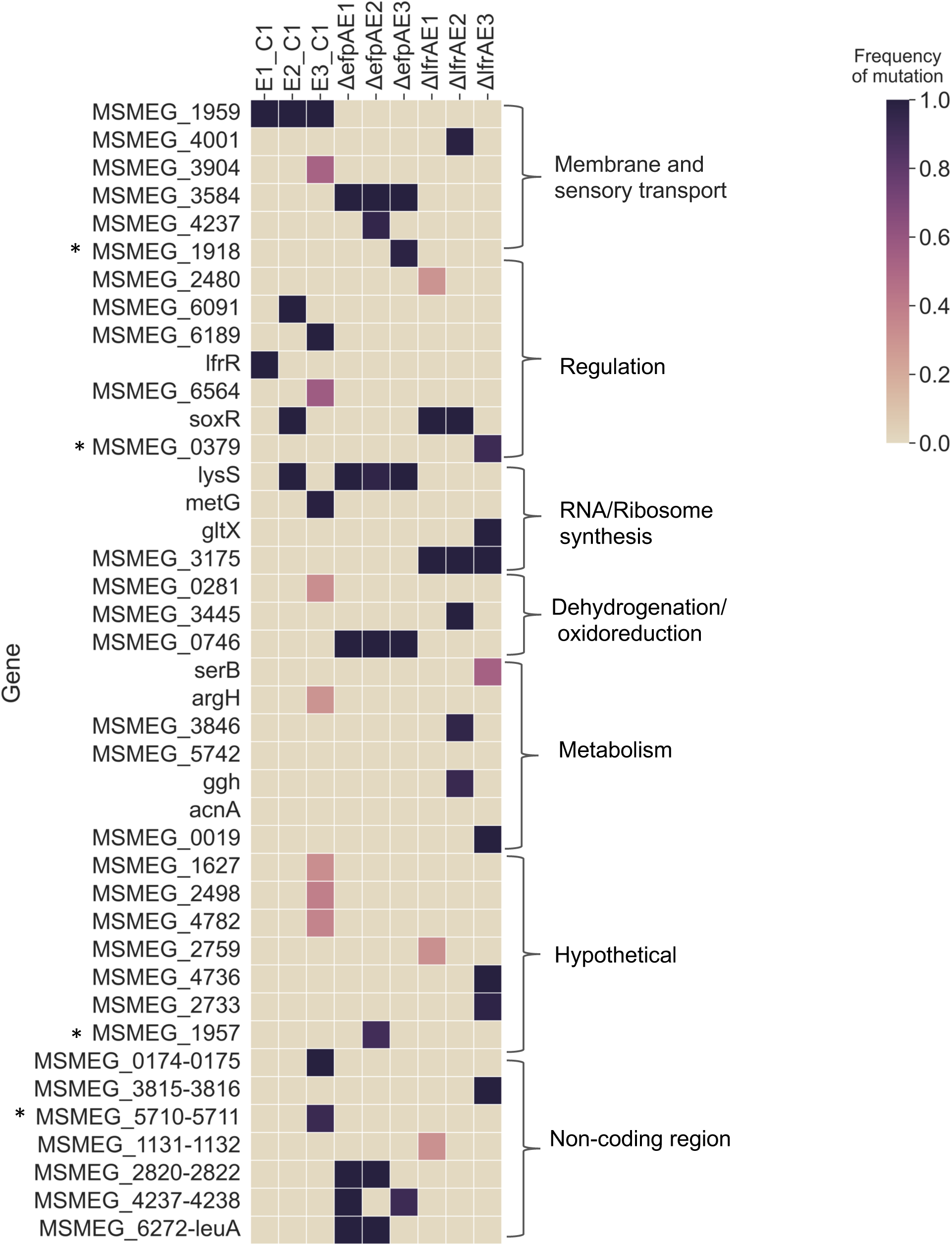
Heatmap of mutations and their frequencies in evolved Msm strains. Heatmap representation of genetic variants identified in wild-type evolved (E1, E2, E3), ΔlfrA evolved (E1, E2, E3), and ΔefpA evolved (E1, E2, E3) cells. Each row corresponds to a specific gene or intergenic region, while each column represents an evolved strain. The colors indicate the frequency of each detected variant, applying a frequency cutoff of > 30% and a minimum coverage depth of 10. Wild-type evolved strains were compared against three WT controls that were evolved without EtBr. The ΔlfrA and ΔefpA evolved strains were compared to their respective controls, evolved without EtBr. All identified variants are single-nucleotide polymorphisms (SNPs), unless marked with an asterisk (*), which indicates insertion or deletion mutations (INDELs). Mutation positions and predicted amino acid changes are provided in Supplementary Data 1.

Interestingly, the Δ*lfrA* evolved strains did not acquire mutations in *MSMEG_1959*, suggesting strain-specific adaptive routes. Instead, all Δ*lfrA* evolved mutants shared a common mutation in *MSMEG_3175*, which encodes a pseudouridine synthase associated with the large ribosomal subunit. Additionally, two of the Δ*lfrA* mutants (E1 and E2) acquired mutations in *soxR*, indicating partial convergence with the wild-type strains in targeting global regulators. The Δ*lfrA*_E3 strain was unique in harboring a mutation in the upstream regulatory region of *MSMEG_3815*, an MFS family efflux pump. This was the only identified mutation within a promoter region of an efflux gene among all evolved populations, suggesting a potential upregulation of this efflux pump in the absence of *lfrA*.

The Δ*efpA* evolved strains (E1, E2, and E3) followed a different evolutionary trajectory. All three lineages accumulated mutations in two membrane protein-encoding genes: *MSMEG_3584*, which belongs to the MmpL family of efflux transporters, and *MSMEG_4237*, encoding a conserved membrane protein with an unknown function. These parallel mutations across all Δ*efpA* lineages suggest a convergent evolutionary path to resistance in the absence of *efpA*. Notably, none of the Δ*efpA* mutants carried mutations in known transcriptional regulators such as *lfrR* or *soxR*, highlighting that the genetic background of the strain strongly influences the specific resistance mechanisms adopted.

Beyond efflux-associated mutations, evolved strains also exhibited changes in genes associated with transcriptional regulation, oxidative processes, dehydrogenation reactions, fatty acid metabolism, and amino acid biosynthesis. These broader mutational patterns suggest that metabolic remodeling accompanies efflux activation, possibly by supporting energy-intensive efflux and mitigating stress from prolonged exposure to EtBr. Together, these genomic data reveal that EtBr resistance in *Msm* can emerge through multiple, background-dependent evolutionary paths that couple efflux activation with membrane and metabolic adaptation.

### RNA-Seq identifies various routes of resistance in mutants

To assess the transcriptional impact of adaptive mutations, we performed whole-transcriptome RNA sequencing (RNA-seq) on evolved wild-type (WT) *Msm* and the efflux pump-deficient strains, Δ*lfrA* and Δ*efpA*. Differential gene expression analysis identified only five differentially expressed genes (DEGs) common across all five evolved strains, including two hypothetical proteins and three enzymes (a flavin-binding monooxygenase, an acyl-CoA synthetase, and a D-amino acid dehydrogenase), highlighting largely strain-specific adaptive responses to EtBr stress (Fig 3A). Within genetic backgrounds, overlap increased. For example, WT_E1 and WT_E2 shared 15 DEGs, including a putative membrane protein (*MSMEG_0172*), oxidoreductases, dehydrogenases, and monooxygenases that were upregulated, whereas sugar ABC transporters were consistently downregulated. In the Δ*lfrA*_E1 and Δ*lfrA*_E3, a larger transcriptional shift was observed, with 1500–1600 DEGs in each strain, of which 465 were shared (Fig 3A).

**Fig 3:**
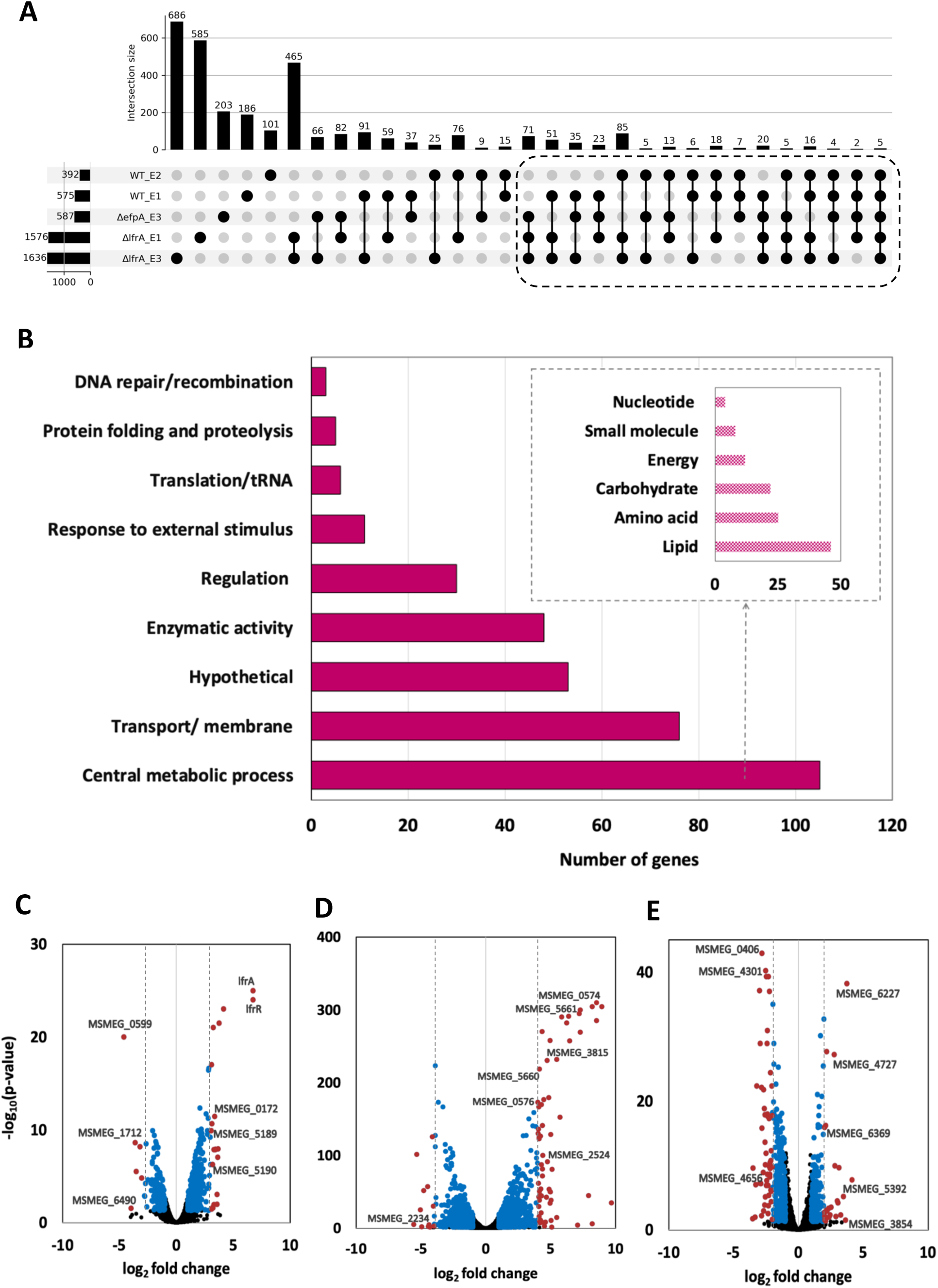
Overview of differentially expressed genes (DEGs) across evolved strains. **(A)** Upset plot illustrating the total number of DEGs (X-axis) across all samples and their intersections across evolved strains (Y-axis). Each column represents an individual evolved strain ( first five columns) or a shared set of DEGs among multiple strains, indicated by connected dots and lines below the X-axis. The number of genes in each set is displayed above the corresponding column, while the shared sample sets are indicated in the graphic below each column, with sample names listed on the left. **(B)** Functional annotation of the 366 common DEGs identified in the upset plot includes sets of genes present in at least three out of the five evolved strains. The DEGs presented have a log_2_ fold change (log_2_FC) greater than ±1 with a significance level of p < 0.05. Genes belonging to other small categories are excluded from the plot and are reported in Supplementary Data S2. **(C-E)** The volcano plots display the log_2_FC of differentially expressed genes across the evolved strains from different genetic backgrounds: WT_E1, ΔlfrA_E1, and ΔefpA_E3, respectively. Genes with a p-value <0.05 and a log_2_FC greater than ±1 are marked in blue. Among these, genes exhibiting a log_2_FC greater than ±3 in WT_E1, greater than ±4 in ΔlfrA_E1, and greater than ±2 in ΔefpA_E3 are highlighted in red. All remaining non-significant genes are represented in black. Plots for WT_E2 and ΔlfrA_E3 are shown in Fig S4.

Across all five strains, 366 DEGs were identified as common to at least three strains, as highlighted in Fig 3A. Functional annotation of these revealed that a significant portion (76 genes) was linked to membrane transport, including various efflux pumps, and 30 were transcriptional regulators (Fig 3B). This supports the idea that membrane remodeling and regulatory rewiring are key to the adaptive response to EtBr stress. In addition to transport, a significant number of genes were involved in central metabolic pathways. Lipid metabolism was the most significantly impacted pathway, with 46 DEGs involved in lipid biosynthesis and breakdown. Other enriched pathways included genes from carbohydrate metabolism (22 DEGs), amino acid metabolism (25 DEGs), and energy metabolism (12 DEGs).

Among the enzymatic DEGs, 48 belonged to essential catalytic families, including dehydrogenases, acetyltransferases, monooxygenases, and methyltransferases. Additional pathway-specific changes were noted in genes involved in responses to external stimuli, protein translation, tRNA modification, protein folding, proteolysis, and DNA repair and recombination. Analysis of the most strongly regulated genes across evolved backgrounds indicates heterogeneous yet coordinated transcriptional responses. In the WT_E1 lineage, a large subset of transporters, regulators, and metabolic genes exhibited expression shifts exceeding ±3 log₂FC (Fig 3C). Prominent among these were the efflux transporters *lfrA* (log₂FC 6.7) and *MSMEG_0172* (log₂FC 3.36), whereas *MSMEG_1712* (log₂FC –3.6) and *MSMEG_6492* (log₂FC –3.9) were significantly downregulated. Regulatory genes, including *lfrR*, *MSMEG_5190*, and the oxidoreductase *MSMEG_0589,* also showed strong upregulation, while *MSMEG_0599*—encoding an acyl-CoA synthetase—was markedly repressed.

In ΔlfrA_E1, loss of *lfrA* was compensated by the induction of multiple alternative efflux-associated genes, including *MSMEG_3815, MSMEG_5660, MSMEG_5661, MSMEG_2524,* and *MSMEG_0576* (Fig 3D). Additionally, sigma factor genes *MSMEG_0573* and *MSMEG_0574 (rpoE family)* were highly upregulated (log₂FC 9.6 & 8.5), accompanied by downregulation of *MSMEG_2234*, an acyl-CoA dehydrogenase.

The ΔefpA_E3 strain displayed a broader but more moderate response relative to WT and ΔlfrA strains. Several efflux pumps exhibited mild upregulation, including the RND transporter *MSMEG_4741* (log₂FC 1.08) and multiple MFS transporters (*MSMEG_6225, MSMEG_6922, MSMEG_5556*, and *MSMEG_4767*, each with a log₂FC of 1.0–1.6). Among the most highly induced genes was *MSMEG_6227*, a PadR-family transcriptional regulator (log₂FC 3.7; Fig 3E), which is known to regulate efflux systems and stress-associated pathways (34, 35). Additional transporters, such as *MSMEG_3854* and *MSMEG_4656,* were upregulated, as was *MSMEG_6369*, an ATP-binding protein exporter (log₂FC 2). Differentially expressed metabolic genes included *acyl-CoA dehydrogenase (MSMEG_0406)*, *MSMEG_4301 synthase*, and *mycocerosic acid synthase (MSMEG_4727)*.

Collectively, these transcriptomic patterns demonstrate that EtBr resistance arises not only from enhanced efflux activity but also through extensive reprogramming of membrane transport, regulatory hierarchies, and central metabolic pathways.

### Distinct molecular routes are employed by the mutants to achieve efflux activation

To explore the relationship between membrane transport rewiring and multidrug resistance potential in the evolved *Msm* strains, we assessed the expression profiles of efflux pumps across all transcriptomic datasets. Transport and membrane genes were compared to annotated multidrug transporters from the TransportDB database (35), which identified 101 multidrug transporters in *Msm*. Of these, 43 showed significant differential expression across the evolved populations (Fig 4A). These efflux pumps belonged to four major transporter superfamilies—ATP-binding cassette (ABC), major facilitator superfamily (MFS), resistance–nodulation–cell division (RND), and multidrug/oligosaccharidyl-lipid/polysaccharide (MOP) flippases (Fig 4B). Notably, MFS transporters constituted the majority, with 31 of 43 DEGs, highlighting their dominant contribution to efflux-mediated resistance under EtBr exposure. RND transporters represented the next major group, consistent with their widely recognized role in intrinsic and acquired drug resistance in mycobacteria. Overall, the Δ*lfrA* evolved populations exhibited a distinct increase in the number of differentially expressed drug transporters. Both Δ*lfrA*_E1 and Δ*lfrA*_E3 showed significant regulation of approximately 30 efflux pumps, compared with only ∼10 pumps in WT_E1, WT_E2, and Δ*efpA*_E3. Notably, efflux modulation in these strains was not uniformly unidirectional; while several pumps were strongly upregulated, others were concurrently downregulated. Of particular interest, efpA—a well-characterized MFS pump—was repressed in three evolved lineages, including both Δ*lfrA* strains, suggesting potential regulatory redistribution or functional compensation between efflux components under EtBr selection (Fig 4A).

**Fig 4:**
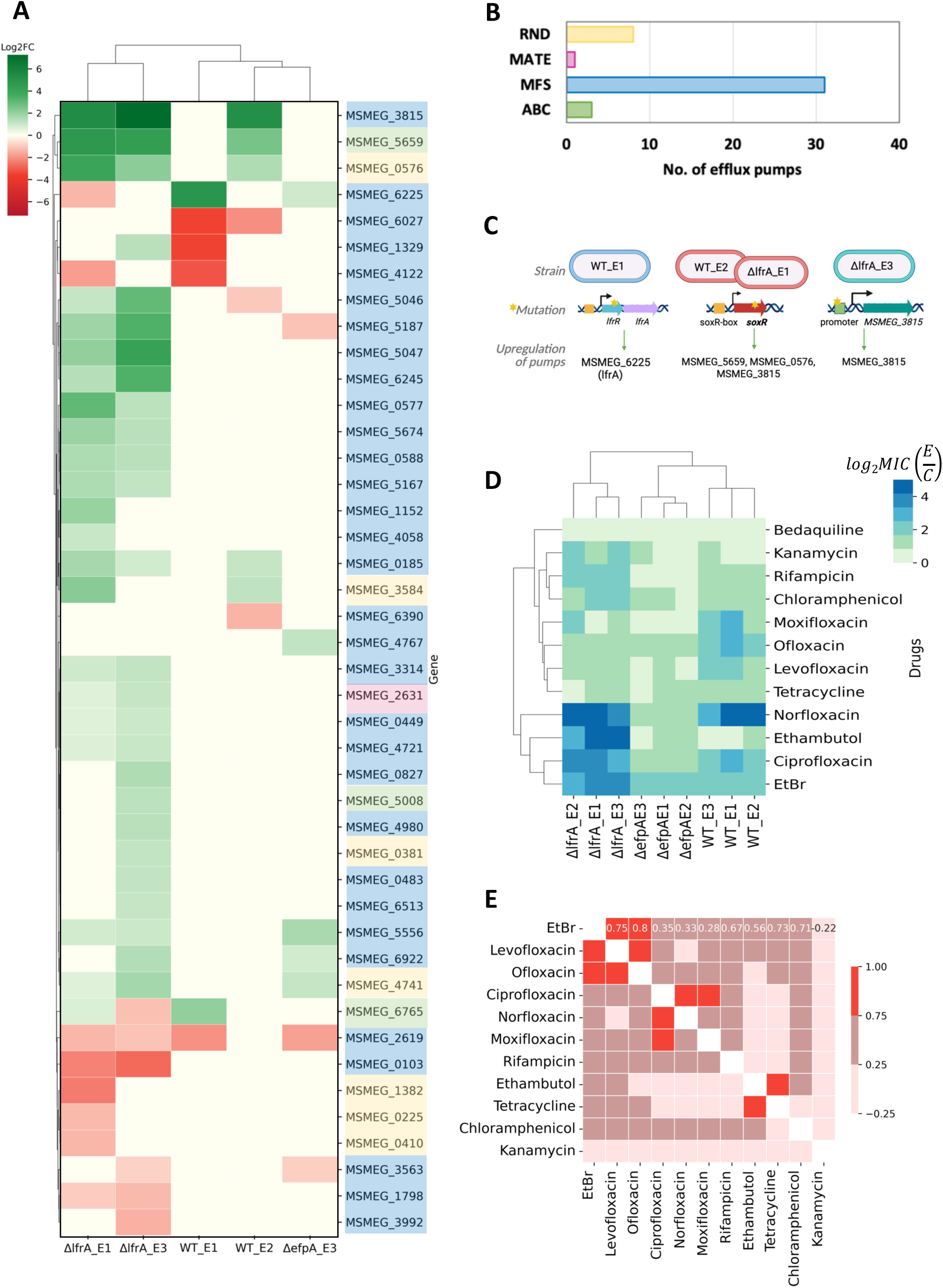
Efflux pump upregulation mediates resistance across all evolved mutants. **(A)** Heatmap of multidrug efflux pump genes differentially expressed in at least one evolved mutant (RNASeq, p<0.05). The heatmap is drawn based on the log2FC values. The color-coded gene name column indicates efflux transporter classes, as defined in B. **(B)** Bar plot showing the distribution of differentially expressed efflux pump genes across various transporter classes. **(C)** Schematic illustrating the relationship between efflux pump expression and the corresponding mutations identified across evolved mutants **(D).** Cross-resistance of the mutants WT E1, E2, E3, ΔlfrAE1, E2, E3, and ΔefpAE1, E2, E3 towards 12 drugs is indicated as log2 fold change MIC increase with respect to their corresponding control strains. (MIC calculated by the broth microdilution method and resazurin assay. n=3 biological replicates). **(E)** Correlation matrix of MIC values of all drugs across all evolved mutants.

Integrating genomic and transcriptomic data revealed that efflux upregulation emerges through distinct yet convergent evolutionary routes (Fig 4C). In WT_E1, an SNP in the repressor *lfrR* (*MSMEG_6223*) enabled high constitutive expression of *lfrA*, driving resistance largely through a single dominant efflux system. By contrast, WT_E2 and ΔlfrA_E1 exhibited activation of multiple pumps, including MSMEG_5661, MSMEG_0576, and MSMEG_3815, likely mediated by mutations in soxR, consistent with its role as a global redox-responsive regulator. In ΔlfrA_E3, strong induction of MSMEG_3815 correlated with an SNP in its promoter region, suggesting promoter-level activation (Fig 4C). Together, these observations demonstrate that efflux intensification can arise through the relief of local repression, global regulatory rewiring, or promoter gain-of-function, each producing a distinct yet effective resistance trajectory. Consistent with these mechanistic variations, WT and ΔlfrA populations primarily achieved resistance through the strong upregulation of efflux driven by regulatory mutations, whereas ΔefpA lineages relied on more modest yet distributed activation of multiple pumps, likely accompanied by additional compensatory adaptations. Despite the diversity of underlying changes, all lineages converged on a comparable resistance ceiling, indicating an upper operational limit to efflux-mediated resistance.

To assess whether enhanced EtBr efflux conferred broader antibiotic resistance, we measured MICs for evolved wild-type, ΔlfrA, and ΔefpA strains across a panel of drugs (Fig 5D). All strains except *ΔefpA* evolved ones exhibited strong cross-resistance to fluoroquinolones, with norfloxacin MICs rising 8–16-fold and ciprofloxacin 4–8-fold in both wild-type and *ΔlfrA* backgrounds. Evolved *ΔlfrA* mutants further displayed marked resistance to rifampicin, suggesting potential co-regulation of rifampicin-associated transport pathways, and showed increased resistance to tetracycline, kanamycin, and chloramphenicol. A particularly notable shift was the 8–16-fold increase in ethambutol resistance in *ΔlfrA* strains, indicative of transcriptional or mutational adaptations altering ethambutol sensitivity. In contrast, *ΔefpA* mutants displayed only a modest increase in MICs, with fluoroquinolone resistance elevating 2–4-fold, consistent with their comparatively weaker efflux induction (1–2 log₂FC).

**Fig 5:**
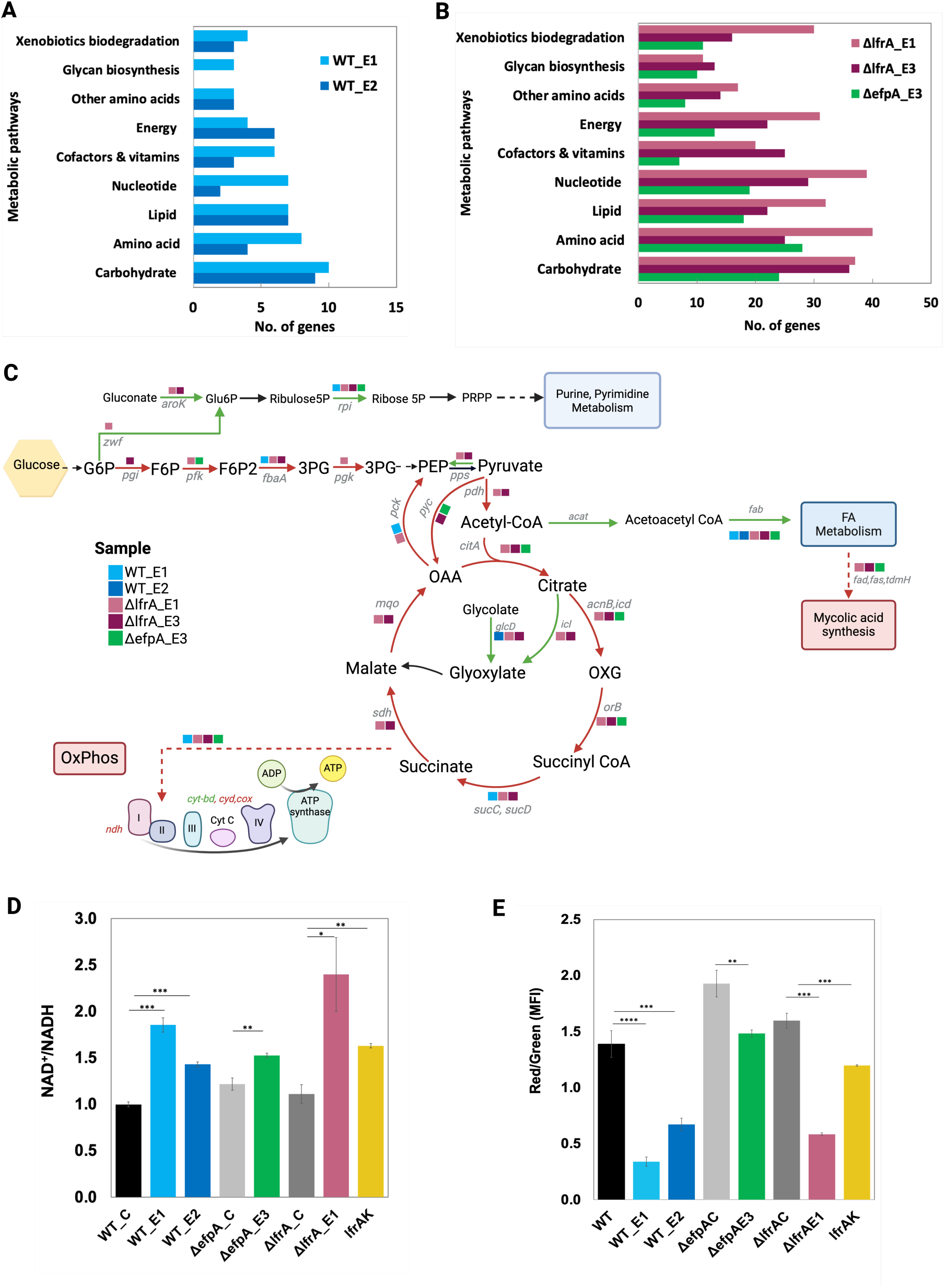
Metabolic shift across evolved mutants. **(A)** Gene enrichment analysis of DEGs in WT_E1 and WT_E2 evolved mutants. **(B)** Gene enrichment analysis of the DEGs in evolved mutants ΔlfrAE1, ΔlfrAE3, and ΔefpA3. KEGG pathways were used for the gene enrichment analysis.**(C)** Metabolic map of Msm illustrating key pathways and enzymes affected in the evolved strains. Green arrows indicate upregulated DEGs, while Red arrows indicate downregulation DEGs. The colored boxes highlight the evolved strains in which these DEGs occur. Log2 fold change values of these genes are provided in Supplementary Data 3. **(D)** Quantification of NAD+/NADH levels in the mutants and their respective controls using a redox cycling assay. **(E)** Measurement of membrane potential using DiOC_2_ and a flow cytometer. The red-to-green fluorescence ratio, indicative of membrane potential, is shown for each strain. * Indicates p-value <0.05, ** indicates p-value <0.01,*** indicates p-value <0.001 and **** indicates p-value <0.0001

Correlation analysis between EtBr MICs and MICs for 12 antibiotics (Fig 4E) further supported this relationship: 11 drugs showed positive association with EtBr resistance, with the strongest correlations observed for levofloxacin (r = 0.75) and ofloxacin (r = 0.80). Moderate correlations with tetracycline, chloramphenicol, ethambutol, and rifampicin underscore broader efflux-mediated resistance beyond EtBr. Together, these findings suggest that EtBr resistance co-occurs with multidrug resistance, consistent with increased efflux activity across evolved strains.

### Metabolic rewiring observed in mutants

To further understand the functional impact of differential gene expression in the evolved *Msm* strains, we performed gene enrichment analysis to identify significantly altered metabolic and regulatory pathways. Across all evolved strains regardless of genetic background, we observed a consistent enrichment of genes involved in carbohydrate and amino acid metabolism, which formed one of the largest functional categories among the differentially expressed genes (Fig 5A & 5B). Additionally, genes associated with energy metabolism, lipid metabolism, nucleotide metabolism, vitamins and cofactors metabolism, and glycan biosynthesis were significantly enriched.

Mapping these differentially expressed genes onto central metabolic pathways (Fig 5C) revealed a pronounced metabolic rewiring. Genes involved in glycolysis were consistently downregulated across all evolved mutants, including key enzymes such as phosphoglucomutase (pgm), glucose phosphate isomerase (pgi), phosphofructokinase (pfk), fructose-bisphosphate aldolase (fbaA), and phosphoglycerate kinase (pgk). Likewise, genes encoding enzymes of the tricarboxylic acid (TCA) cycle—including citrate synthase (citA), isocitrate dehydrogenase (icd1), oxoglutarate dehydrogenase (orB), succinyl-CoA synthetase (suc), and succinate dehydrogenase (sdh)—were also downregulated, indicating a broad suppression of central carbon metabolism. This downregulation likely limits NADH production from conventional carbon oxidation routes.

In contrast, genes of the pentose phosphate pathway, especially ribose-5-phosphate isomerase (rpi), were upregulated, suggesting a redirection of carbon flux toward biosynthesis and NADPH generation, thus favoring anabolic growth and redox homeostasis over ATP production. Notably, there was upregulation of a FAD-binding glycolate dehydrogenase (glcD) and isocitrate lyase (icl), strongly implicating activation of the glyoxylate shunt as an alternative carbon-conserving and redox-modulating pathway. Glycolate dehydrogenase likely contributes to FADH₂-driven electron input into the ETC, partially compensating for reduced NADH levels, while isocitrate lyase bypasses NADH-generating steps of the TCA, supplying glyoxylate and succinate to sustain biomass production under metabolic constraint.

Fatty acid metabolism genes exhibited a mixed transcriptional profile (Fig. 5C and Supplementary Data 3); however, several genes associated with β-oxidation were upregulated, suggesting that the catabolism of endogenous or exogenous lipids served as an alternative energy source. This is consistent with increased reliance on acetyl-CoA generation via lipid degradation, which could feed into the glyoxylate cycle and support residual oxidative phosphorylation.

Importantly, the downregulation of TCA enzymes was paralleled by a reduction in the expression of oxidative phosphorylation genes, such as succinate dehydrogenase (sdh) and cytochrome oxidase (cydB). However, WT_E1 and WT_E2 strains showed a modest (log₂ 1–2 fold) upregulation of the cytochrome bd oxidase (cyt-bd), a high-affinity terminal oxidase that functions under oxygen-limited or redox-impaired conditions. This suggests a shift away from classical aerobic respiration toward an ETC configuration favoring limited oxygen utilization and redox flexibility. Such remodeling, along with diminished NADH production, indicates a metabolic state adapted for redox balance rather than maximal energy output. To validate this hypothesis, we quantified NAD⁺ and NADH levels using a redox cycling assay. Evolved strains exhibited a ∼2-fold reduction in NADH levels as compared to their controls, leading to a significantly increased NAD⁺/NADH ratio (Fig 5D), confirming a redox state skewed toward oxidation.

In addition to this, the upregulation of PMF-dependent efflux pumps would further drain the proton motive force, potentially contributing to membrane depolarization. Since metabolically active bacterial cells maintain a membrane potential (MP) of approximately –150 mV, largely driven by ion gradients established via the ETC, any disruption in electron flow or proton pumping could collapse this potential. Using the DiOC₂ (3) dye assay, we observed a significant reduction in membrane potential across all resistant strains—WT_E1, WT_E2, ΔlfrA_E1, and ΔefpA_E3—compared to their respective controls (Fig 5E). Notably, the LfrAK strain, which overexpresses the LfrA efflux pump, also displayed a reduced membrane potential, further supporting the idea that efflux-driven PMF consumption, coupled with altered electron input and oxygen limitation, underlies the observed depolarization. Thus, these findings suggest that metabolically rewired, efflux-overexpressing strains adopt a survival-oriented metabolic state characterized by suppressed NADH-dependent respiration, redox-compensating pathways like the glyoxylate shunt, partial reliance on FAD-linked electron donors, and a shift to low-energy ETC configurations, ultimately leading to a reduction in both energy generation and membrane potential.

Interestingly, despite EtBr being a DNA intercalating agent known to trigger nucleic acid damage, SOS response or DNA repair pathways were not enriched in any of the evolved strains, likely because RNA-Seq was performed on adapted strains in the absence of acute EtBr stress.

Overall, gene enrichment and pathway analyses reveal that the evolved *Msm* strains have undergone extensive metabolic reprogramming, including suppression of central carbon metabolism, increased expression of membrane transporters and efflux pumps, and disruption of redox balance and membrane potential. These changes likely contribute in concert to the high-level EtBr resistance and multidrug cross-resistance.

### Limit of efflux-mediated resistance in *M tuberculosis* H37Ra

We further extended our study to *Mtb* by evolving the avirulent H37Ra strain under progressively increasing concentrations of EtBr. Similar to *Msm*, all evolved *Mtb* lineages exhibited a resistance plateau, adapting to up to 8 times the MIC of EtBr. These evolved populations exhibited a 16-fold increase in EtBr MIC, accompanied by a reduction in intracellular EtBr accumulation (Fig 6A & B). Whole-genome sequencing revealed mutations in genes encoding a transcriptional regulator, MCE family proteins, PE/PPE proteins, and several others (Fig 6D). Notably, mutations in the transcriptional regulator MRA_3099 were consistently observed across all three evolved lineages, suggesting potential upregulation of the adjacent mmr efflux pump gene (MRA_3098). Transcriptomic profiling of the *Mtb_E3* evolved strain confirmed a significant upregulation of the mmr efflux pump (MRA_3098), with a log₂ fold change of 3.6 (Fig 6E). Additionally, 11 other multidrug efflux pump genes were upregulated among the differentially expressed genes (Fig 6E), reinforcing the enhanced efflux activity in the evolved strain. Consistent with observations in *Msm*, this increase in efflux activity was associated with a reduction in membrane potential (Fig 6C). Metabolic profiling further revealed metabolic rewiring in the evolved *Mtb* populations, mirroring the adaptations observed in *Msm* (Fig 6F). Genes involved in glycolysis and the oxidative branch of the TCA cycle were downregulated, whereas those associated with gluconeogenesis and the glyoxylate shunt were upregulated. Moreover, multiple genes linked to fatty acid metabolism exhibited both upregulation and downregulation, indicating a broader reorganization of carbon flux and energy metabolism in response to EtBr selection pressure.

**Fig 6:**
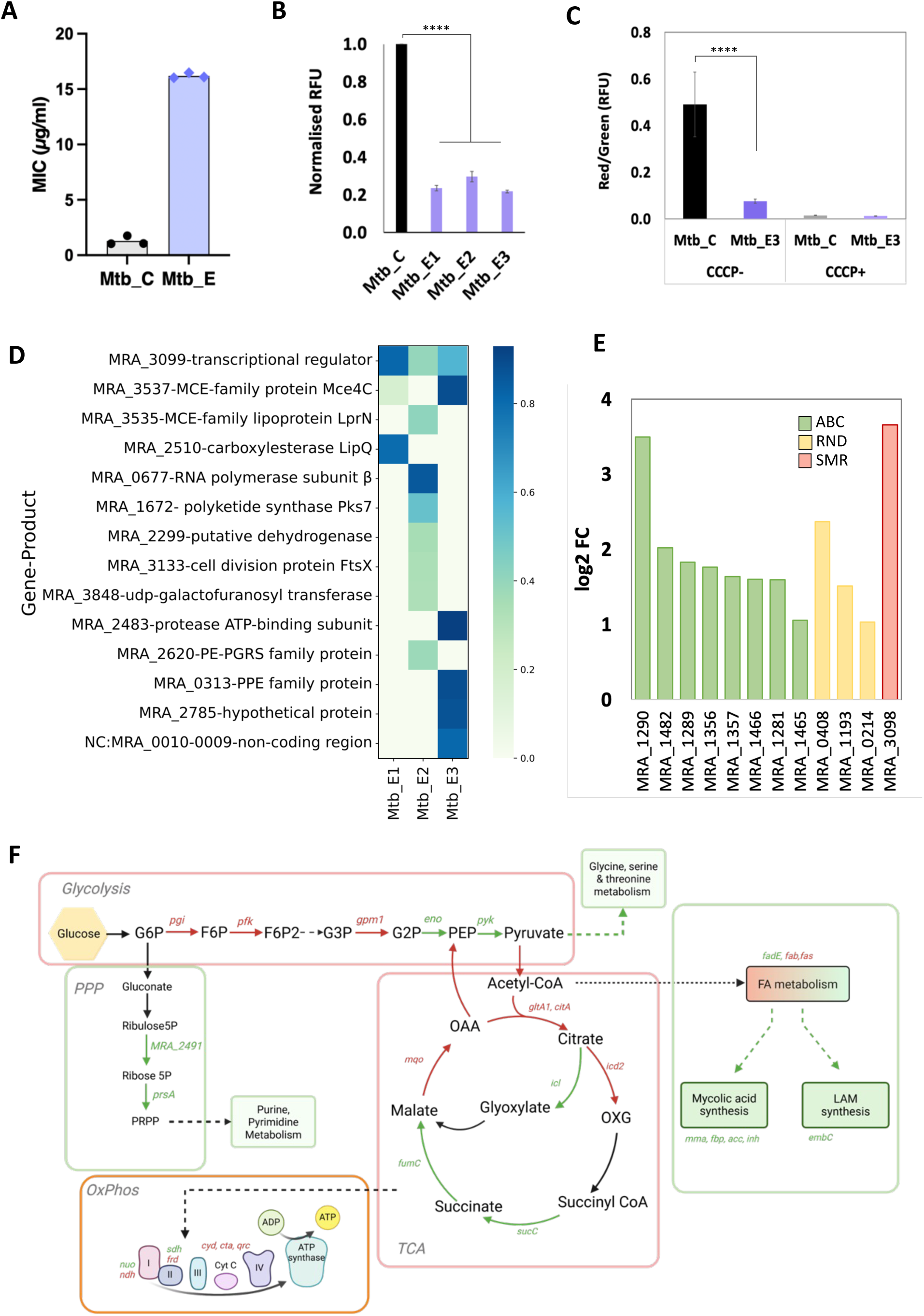
Testing the limit of efflux resistance in Mtb. **(A)** EtBr MIC for the evolved Mtb lineages (E1, E2, and E3), and Mtb_C, the control strain evolved in the absence of EtBr.**(B)** EtBr uptake in control and mutant strains at an EtBr concentration of 1μg/ml, and accumulation measured for 1 hour. End-point readings normalised to control are plotted here. **(C)** Membrane potential measured using DiOC_2_. The red-to-green fluorescence ratio, indicative of membrane potential, is shown for the Mtb_C and Mtb_E3 evolved strain. CCCP at a concentration of 10ug/ml is used to disrupt the membrane potential. **(D)** Heatmap indicating the gene mutations and their frequencies in the Mtb evolved (E1, E2, E3) populations. Frequency greater than or equal to 30% and coverage depth cutoff of 10 is considered. The mutation position and type of change in the amino acid are mentioned in Supplementary Data 1. **(E)** Gene expression indicated as log_2_ fold change of the differentially expressed multidrug efflux pumps in Mtb_E3. **(F)** Metabolic map illustrating key metabolic pathways in Mtb. DEGs that are upregulated are indicated in green, whereas the ones that are downregulated are in red. Log2 fold change values of these genes are provided in Supplementary Data 4. **** indicates p-value <0.0001

## Discussion

Antimicrobial resistance represents a growing global health threat, contributing to millions of deaths each year. Among the various mechanisms that bacteria employ to evade antibiotics, drug efflux has emerged as a central contributor to multidrug resistance. While previous studies have demonstrated the role of efflux pumps in mediating resistance, they have not clearly defined the upper threshold of resistance that can be achieved through efflux pump overexpression alone. Our study addresses this gap by employing a broad-spectrum efflux substrate, EtBr, in an adaptive laboratory evolution framework to systematically investigate the limits of efflux-based resistance. EtBr, a DNA intercalating agent and a known substrate for several multidrug efflux pumps, was used to selectively enrich for mutations associated with efflux activity. Through stepwise exposure to increasing EtBr concentrations, we observed that all evolved *Msm* and *Mtb* populations were able to adapt up to 4-16× their minimum inhibitory concentration (MIC), but no lineage was capable of surpassing this threshold—even under extended selection pressure. This plateau suggests an upper bound to resistance that can be achieved solely through efflux pump activation in these systems.

Whole-genome sequencing revealed that adaptation to EtBr was largely driven by mutations in regulatory genes, rather than structural alterations in efflux pump proteins themselves. This observation aligns with a well-established trend in clinical isolates, where regulatory gene mutations—for example, marR in *Escherichia coli* (36), mepR in *Staphylococcus aureus* (37), mexR in *Pseudomonas aeruginosa* (38), and acrR and ramA in *Klebsiella pneumoniae* (39)—are widely reported. In *Mtb*, recurrent mutations in Rv0678, the regulator of the MmpL5 efflux pump, confer resistance to bedaquiline and are prevalent in clinical strains (40). Structural mutations in efflux pumps are less frequently observed, though some clinical cases implicate changes in AcrB (*Escherichia coli* and *Salmonella enterica*) (41,42), Tap (*Mtb*) (43), MtrD (*Neisseria gonorrhoeae*) (44), and AdeJ (*Acinetobacter baumannii*) (45). Consistent with these patterns, our study identified mutations exclusively in local regulators, such as lfrR (the repressor of lfrA in *Msm*), MRA_3099 (the repressor of mmr in *Mtb*), and in global transcriptional regulators, notably soxR. Importantly, we did not detect any structural mutations within efflux pump genes in our evolved populations. As a redox-sensitive global regulator, *soxR* controls genes that respond to oxidative stress, and in our study, these mutations correlated with the upregulation of multiple efflux pumps as well as sigma factors involved in stress adaptation. These findings parallel earlier reports in *E. coli* and *Salmonella enterica*, where *soxR* mutations enhanced resistance through combined activation of oxidative stress defenses and efflux pathways (46,47). Our work extends this understanding to *Msm*, where the role of *soxR* in AMR had not been well characterized.

Genetic background influenced mutation trajectories, highlighting potential epistatic effects. Our whole-genome sequencing data revealed notable genotype-specific mutation patterns that reflect the influence of genetic background on the evolutionary trajectory of resistance. Specifically, mutations in the gene *MSMEG_1959*, encoding a putative membrane protein, occurred exclusively in *Msm* wild-type (*WT*) evolved lineages, while regulatory gene mutations, including those in *lfrR* and *soxR*, were restricted to *WT* and Δ*lfrA* strains, and absent in Δ*efpA* strains. This divergence suggests a potential epistatic influence where the presence of specific efflux pumps constrains or guides the acquisition of specific mutations during the evolution of resistance.

Transcriptomic analysis revealed that both *Msm* and *Mtb* evolved strains upregulated either single or multiple efflux pumps to reach a similar resistance plateau. In *Msm*, the pumps that were upregulated were associated with broad cross-resistance, particularly to fluoroquinolones. In contrast, Mtb-evolved cells displayed only moderate cross-resistance, with a ∼2-fold increase in isoniazid MIC (data not shown), likely reflecting the narrower substrate specificity of the induced pumps. Notably, the lfrA pump in *Msm* and the mmr pump in *Mtb* are both known to export quaternary ammonium compounds such as ethidium bromide (48–50). Thus, maximal induction of these pumps alone was sufficient for the WT_E1 Msm population and the Mtb_E3 lineage to reach their respective resistance ceilings. In addition to efflux transporters and their regulators, we observed differential expression of numerous membrane-associated genes and extensive metabolic reprogramming. Notably, genes involved in the TCA cycle, including *citrate synthase*, *isocitrate dehydrogenase*, and *succinate dehydrogenase,* were consistently downregulated. Previous studies in *E. coli* have shown that suppression of aerobic respiration can increase antimicrobial resistance (51). In our evolved populations, this metabolic shift was accompanied by an increased reliance on the glyoxylate shunt and pentose phosphate pathway, a strategy also observed in *Mtb* during intracellular survival (52,53). These metabolic alterations resulted in reduced intracellular NADH levels and a higher NAD⁺/NADH ratio, a redox state previously linked to improved persistence in *Mtb* (54). A similar artificially induced increase in NAD⁺/NADH has been shown to enhance survival in *Pseudomonas aeruginosa* (55). At the same time, the overexpression of energy-dependent efflux pumps disrupted the PMF, leading to measurable membrane depolarization. Membrane potential assays confirmed, for the first time in *Msm and Mtb*, that resistant strains exhibited lower PMF than their parental control. This finding is consistent with studies in *Salmonella typhimurium*, where the deletion of efflux pumps increased PMF, implicating active efflux in the dissipation of PMF (56).

Together, our findings demonstrate that the evolution of efflux-mediated resistance is shaped not only by the selection of specific efflux pumps and regulatory mutations but also by broader cellular adaptations involving metabolic and redox homeostasis. The interplay between genetic background, regulatory network reprogramming, and metabolic shifts defines the resistance landscape under selective pressure. These insights underscore the multifaceted nature of antimicrobial resistance evolution and highlight the importance of considering both efflux-dependent and efflux-independent mechanisms when developing strategies to counteract AMR.

## Materials and Methods

### Bacterial strains and media

In the evolution experiments, the *Mycobacterium smegmatis mc2* wild-type strain, along with the efflux pump-deleted strains *ΔefpA* and *ΔlfrA, and Mycobacterium tuberculosis* H37Ra, were utilized. The *ΔlfrA* strain was received from Prof. Miguel Viveiros, GHTM, Lisboa, Portugal, as a kind gift. Knockout strain of MSMEG_2619 (*ΔefpA*) gene was obtained from Todd Gray, NYSDOH Wadsworth Center, Albany, New York. All strains (wild type and evolved mutants) were cultured in Middlebrook 7H9 broth (HiMedia) supplemented with 0.44% (v/v) 100% glycerol and 0.15% (v/v) Tween 80 at 37 °C in 200 rpm shaking conditions. For *Mtb* strains, 10% of ADC supplement was added along with M7H9. For plating the cells, Luria Bertani (HiMedia) agar plates were used. EtBr, Chloramphenicol, Rifampicin, Kanamycin, Tetracycline, and Ethambutol were sourced from HiMedia. Fluoroquinolones Norfloxacin, Ciprofloxacin, Levofloxacin, Ofloxacin, and Moxifloxacin were from Sigma.

### Evolution experiment

*Msm* WT, *ΔlfrA, ΔefpA,* and *Mtb* strains were cultured to an OD of 0.5, after which approximately 10⁶ cells were inoculated into 5 mL M7H9 broth containing 0.25× MIC of ethidium bromide (EtBr; HiMedia). Parallel control cultures without EtBr were inoculated with an equivalent number of cells. For all strains, three independent lineages each were propagated. Cultures were incubated at 37 °C with shaking conditions until they again reached an OD of 0.5, at which point they were passaged into fresh medium containing a twofold higher concentration of EtBr. At every passage, 2 mL of culture was stored as glycerol stocks at –80 °C. This stepwise evolution was continued until the populations failed to reach an OD of 0.5 at a given EtBr concentration, at which point the experiment was terminated. For all the *Msm* strains, the evolved populations that tolerated the highest EtBr concentrations were subsequently grown without EtBr and streaked to isolate single colonies for downstream characterization. For *Mtb*, populations from the final passage were revived in EtBr-free medium and similarly characterized.

### Antimicrobial susceptibility assay

#### Broth Microdilution method

MIC determination was performed using a broth microdilution assay in 96-well transparent, sterile microplates (Thermo Fisher Scientific, USA). In each well of the microplate, 200 μL of a 10^5^ cells/mL suspension (corresponding to an OD of 0.005) was added. To each well, the desired antibiotic concentration was added along with the desired concentration of EPI (CCCP) wherever necessary, followed by incubation at 37°C for 48 hours. Post incubation, five drops of 10 μL each from all the wells were dropped on LB agar plates, which were further incubated for 48 hours at 37°C (Drop Count Method), and colony count was done using the formula below:

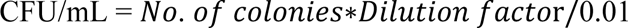

The minimum drug concentration that gave a CFU count of 10^5^ cells/ml or below was considered to be the Minimum Inhibitory Concentration (MIC).

#### Resazurin Microtiter Assay (REMA)

MIC for cross-resistance was detected using the REMA plate assay (57,58). Here again, the cells were grown till mid-log phase, that is, 0.5 O.D. These cells were diluted to obtain 10^5^ cells/mL, of which 200 µL was dispensed in each well of the 96-well plate. The required concentration of the drugs was added in each well. The plates were then incubated for 2 days in the case of *Msm* and 7 days in the case of *Mtb* at 37°C. Post incubation, 10 µL of the culture from each well was plated on Middlebrook 7H11 agar plates. This was done to identify the concentration of the drugs that induced complete sterilization of the culture. The concentration inducing complete clearance is also defined as the minimum bactericidal concentration (MBC). Furthermore, in each well, 60 µL of 0.02% v/v resazurin was added. This setup was again incubated at 37°C for 4 hours (Msm) to 1 day (Mtb) to complete the conversion of resazurin to resorufin. The lowest concentration at which the blue colour of resazurin was sustained after incubation was identified as the MIC.

### EtBr accumulation assay

The EtBr accumulation assay was performed according to the described protocols (32). *Msm* and *Mtb* cells (WT and mutants) were grown in the M7H9 media till they reached an OD of 0.5. After centrifugation at 12,500 rpm for 10 minutes to wash off the media, the cells were resuspended in 1 mL of 1X chilled PBS. Sub-MIC concentrations of EtBr were added to the cells. For some assays, CCCP at a concentration of 10μg/ml was added along with EtBr to inhibit efflux. The intracellular accumulation was determined over a 60-minute period by measuring fluorescence at an excitation wavelength of 530 nm and an emission wavelength of 585 nm using a multiplate reader (Molecular Devices). The accumulation was studied at 37 °C to allow for maximum diffusion of EtBr, considering negligible fluorescence arising from cell adherence. EtBr added to PBS was taken as the negative control for the assay.

### EtBr efflux assay

The EtBr efflux assay was modified as described in earlier protocols (59). *Msm* cells (WT and mutants) were grown in the M7H9 media till 0.5 O.D. After centrifugation at 12,500 rpm for 10 minutes to wash off the media, the cells were resuspended in 1 mL of chilled PBS. To this, EtBr at a subinhibitory concentration of 3.5μg/ml was added along with 10μg/ml of CCCP to allow maximum accumulation of EtBr and limit efflux. The cells were then incubated for 1 hour at RT. After this, the cells were washed once with PBS and then resuspended in 1 mL of chilled PBS. Efflux was measured for 60 minutes at 37 °C by measuring fluorescence at an excitation wavelength of 530 nm and an emission wavelength of 585 nm using a multiplate reader (Molecular Devices).

### DNA extraction and Whole Genome Sequencing

*Msm* and *Mtb* control and evolved mutants were revived from the glycerol stocks and grown in M7H9 broth till mid-log phase, followed by DNA extraction as previously described (60). The DNA was dissolved in 50 µL of TE buffer and stored at -20°C. RNase (Thermo Fisher Scientific) at a concentration of 50 μg/ml was added to the extracted DNA to remove the RNA contamination. The pure DNA was quantified using a Qubit fluorometer (Thermo Fisher Scientific), and outsourced for paired-end sequencing to Novogene Co., Ltd, and HaystackAnalytics Private Limited using the Illumina HiSeq platform with a read length of 150 bp. For some samples, in-house DNA library preparation was carried out using Nextera XT library preparation kit (Illumina). Further QC of the DNA libraries was done using the 4150 TapeStation System (Agilent), and sequencing was carried out using the NextSeq 550 Illumina sequencer with a read length of 75bp.

#### WGS data analysis

Analysis was conducted on the Galaxy platform. The quality of the data was first assessed using FastQC (61). Then, the sequences were mapped to the reference genomes of *Msm* (NC_008596.1) and *Mtb* (NC_009525.1) using the Bowtie2 tool (62). Duplicate reads were removed using the Mark Duplicates tool (63). For variant detection (both SNPs and INDELs), the VarScan(64) tool was employed with the following parameters: minimum coverage depth of 8X, minimum variant frequency of ≥0.01, minimum average base quality of ≥20, and a Phred score of ≥Q20 (>99% accuracy). A coverage depth cutoff of 10 and a variant frequency cutoff of 30% were used. To identify unique mutations, only those absent in all control populations and present in the evolved populations—regardless of depth and frequency cutoffs—were considered. All the *Msm* wild-type evolved strains were compared to three wild-type controls evolved without drugs. *ΔefpA*, *ΔlfrA,* and *Mtb* evolved strains were similarly compared to their corresponding evolved controls. The final mutations were checked by visualizing them using IGV. Heatmaps were generated in Python using Matplotlib v3.5 (65), NumPy v*1.23* (66), Seaborn v0.13 (67) and Pandas v1.4 (68) packages.

#### RNA extraction and library preparation for RNASeq

RNA was extracted from *Msm* and *Mtb* WT and mutant cells, which were grown to mid-log phase and then centrifuged at 4 °C at 8000 rpm for 10 minutes. The cell pellet was stored at -80°C until further processing. The samples were freeze-thawed, and RNA was extracted using TRI reagent (Sigma-Aldrich) according to the manufacturer’s protocol, with slight modifications for lysis. The pellet was suspended in 1 mL TRI reagent, along with 400 µL of 0.1 mm zirconium beads. Cells were mechanically lysed using a bead beater (Neunation iRUPT JR) for 3 cycles (4000 rpm for 1 min; 1 min rest on ice between cycles). DNase (Invitrogen) treatment was applied to all samples for 1 hour at 37°C. The extracted RNA was quantified using a Qubit fluorometer (Thermo Fisher Scientific), and the RNA integrity was determined using a 4150 TapeStation System (Agilent Technologies). RNA with RIN value ≥ 8 were processed for the library preparation. The rRNA was depleted using either Ribo-Zero Plus (Illumina) or NEBNext® rRNA Depletion Kit (New England Biolabs). Finally, the cDNA was synthesized using the Illumina Stranded Total RNA Prep Ligation Kit (Illumina) and the NEBNext Ultra II RNA Library Prep Kit (New England Biolabs). The obtained libraries were quantified using a Qubit fluorometer (Thermo Fisher Scientific) and a 4150 TapeStation System (Agilent Technologies). The validated libraries were outsourced for sequencing to MedGenome Labs Ltd. At least 5 million reads per sample of data were acquired. The raw FASTQ files were analysed on the Galaxy platform (usegalaxy.org) (69).

#### Differential gene expression analysis

The raw FASTQ reads were quality-checked using FastQC (61). The forward and reverse reads from the samples were aligned to the reference genomes using the bowtie2 tool. FeatureCount (70) was used to map the aligned reads to the genes, with the strand parameter set as unstranded and multi-mapping and multi-overlapping features enabled. The read count output from the control and evolved strains was used to identify differentially expressed genes using DESeq2 (71). The significantly (padj<0.05) differentially expressed genes (DEGs) were further processed for gene annotation and functional analysis using the KEGG (Kyoto Encyclopedia of Genes and Genomes) database (72). Gene enrichment was performed using the clusterProfiler (73) and KEGGEnrich packages in RStudio (74). For the analysis of multidrug efflux pumps, the TransportDB (35) database was used to manually curate the list of MDR efflux pumps in *Msm* and *Mtb.* Significant DEGs were then mapped to this list. Gene clustering and visualization were performed in Python using Matplotlib v3.5 (65), NumPy v*1.23* (66), Seaborn v0.13 (67), and Pandas v1.4 (68).

#### NAD^+^/NADH estimation

*Msm* strains were grown to log phase (OD = 0.6), and pyridine nucleotides (NAD^+^/NADH) levels were determined by a redox-cycling assay (75,76). Briefly, 10 mL of each culture was harvested, washed once with 1X PBS, and treated with 0.2 M HCl (1 mL, for NAD+ extraction) or 0.2 M NaOH (1 mL, for NADH extraction). The samples were heated to 55 °C for 20 minutes, followed by cooling to 0 °C for 5 minutes. Samples were then neutralized with 0.1 M NaOH (1 mL, NAD+ extraction) or 0.1 M HCl (1 mL, NADH extraction). After centrifugation, the supernatant containing pyridine nucleotides was passed through a 0.2 µm filter. The redox cycling assay was performed using a reagent mixture consisting of equal volumes of 1 M bicine buffer (pH8.0) (bicine: Sigma Aldrich), absolute ethanol as substrate, 40 mM EDTA (pH8.0), 4.2 mM MTT (3-[4,5-dimethyl thiazol-2-yl]–2,5-diphenyl tetrazolium bromide; thiazolyl blue), and twice the volume for 16.6 mM PES (phenazine ethosulfate; Sigma Aldrich), previously incubated at 30 °C. Of the reagent mixture, 100 μL was incubated with 90 μL of cell extract, followed by the addition of 10 μL alcohol dehydrogenase in 0.1 M bicine buffer, ≈ 6 U for NAD(H) estimation (Sigma Aldrich). The absorbance at 565 nm was recorded every minute for 10 min at 30 °C. Using 0.01–0.1 mM standard solutions of NADH (NADH-Sigma Aldrich) and NAD^+^(NAD- Sigma Aldrich) a standard curve (absorbance versus time plot) was generated whose slope (ΔAbsorbance/ min) of the linear region was correlated to the concentration of coenzyme (mM) by a linear fit equation. This equation was used to determine the concentration(mM) of NAD^+^/NADH in the samples.

### Estimation of membrane potential

The method is adapted from (77), which describes the estimation of membrane potential in Mycobacteria using flow cytometry. Strains were grown in 5ml of M7H9 minimal medium at 37°C. After the cells reached a density of 0.5, they were harvested by centrifugation (8000× g for 10 min), washed once, and then resuspended in sterile PBS. Cells were diluted to approximately 1 × 10^^6^ cells per mL in sterile, filtered PBS. 500 μL of the bacterial suspension was aliquoted into a flow cytometry tube for each staining experiment. To all samples, 3 μL of 3 mM DiOC_2_ was added in each flow cytometry tube and mixed. The samples were incubated at room temperature for 30 min. Samples were then analyzed using a BD FACS Aria Fusion (BD Biosciences) and CFlow Plus software. Data rate was 20000 events/s, and a FSC-H threshold of 8,000 was used to discriminate noise from bacteria. Bacteria were gated using forward scatter and side scatter measurements from blue laser illumination (488 nm), and fluorescence was measured in red and green channels. The ratio of the red and green mean fluorescence intensities (MFI) was plotted for all the samples. In the case of *Mtb* cells, a similar protocol was followed, and fluorescence was measured at 520 nm and 620 nm using the multiplate reader (Agilent BioTek Synergy H1), followed by plotting the ratios of red and green fluorescence.

### Statistical Analysis

A two-tailed Student’s t-test was used to determine the statistical significance with at least three biological replicates. Data with * represents p-value <0.05, ** represents p-value <0.01, *** represents p-value <0.001, and **** represents p-value <0.0001

## Data Availability Statement

The raw genome sequences and RNASeq data for all the strains have been deposited at the NCBI database (BioProject ID: PRJNA1364808)

## Supplementary Information

**Supplementary Data S1** consists of the details of all mutations observed in WGS. Functional category annotation of the DEGs is listed in **Supplementary Data S2.** The DEGs involved in metabolism for *Msm* and *Mtb* are listed in **Supplementary Data S3 and S4,** respectively. The **Supplementary information** file consists of **Supplementary Figures S1, S2, S3, and S4.**

## Author Contributions

**Shiksha Sharma:** Conceptualization, Methodology, Validation, Formal analysis, Investigation, Data curation, Writing - original draft, Visualization. **Sarika Mehra:** Conceptualization, Methodology, Formal analysis, Resources, Data curation, Writing – review & editing, Visualization, Supervision, Project administration, Funding acquisition.

## Acknowledgements

The work was funded by the Department of Science and Technology (DST), India (SERB No. EMR/2016/007667). We thank the Next Generation Sequencing Platform (IoE), Bio-safety Level 2 Facility, and FACS Facility at IIT Bombay. S.S. is supported by the Prime Minister’s Research Fellowship (PMRF ID 1301177)

## References

1. Okeke IN, Kraker MEA de, Boeckel TPV, Kumar CK, Schmitt H, Gales AC, et al. The scope of the antimicrobial resistance challenge. The Lancet. 2024 June 1;403(10442):2426–38.

2. Murray CJ, Ikuta KS, Sharara F, Swetschinski L, Robles Aguilar G, Gray A, et al. Global burden of bacterial antimicrobial resistance in 2019: a systematic analysis. The Lancet. 2022 Feb;399(10325):629–55.

3. Wilson DN, Hauryliuk V, Atkinson GC, O’Neill AJ. Target protection as a key antibiotic resistance mechanism. Nat Rev Microbiol. 2020 Nov;18(11):637–48.

4. Antibiotic Resistance by Enzyme Inactivation: From Mechanisms to Solutions - De Pascale - 2010 - ChemBioChem - Wiley Online Library [Internet]. [cited 2025 June 17]. Available from: https://chemistry-europe.onlinelibrary.wiley.com/doi/full/10.1002/cbic.201000067

5. Wright GD. Bacterial resistance to antibiotics: enzymatic degradation and modification. Adv Drug Deliv Rev. 2005 July 29;57(10):1451–70.

6. Fernández L, Hancock REW. Adaptive and mutational resistance: role of porins and efflux pumps in drug resistance. Clin Microbiol Rev. 2012 Oct;25(4):661–81.

7. Webber MA, Piddock LJV. The importance of efflux pumps in bacterial antibiotic resistance. J Antimicrob Chemother. 2003 Jan;51(1):9–11.

8. Darby EM, Trampari E, Siasat P, Gaya MS, Alav I, Webber MA, et al. Molecular mechanisms of antibiotic resistance revisited. Nat Rev Microbiol. 2023 May;21(5):280–95.

9. Narang A, Garima K, Porwal S, Bhandekar A, Shrivastava K, Giri A, et al. Potential impact of efflux pump genes in mediating rifampicin resistance in clinical isolates of Mycobacterium tuberculosis from India. PLOS ONE. 2019 Sept 26;14(9):e0223163.

10. Consortium TCr. A data compendium associating the genomes of 12,289 Mycobacterium tuberculosis isolates with quantitative resistance phenotypes to 13 antibiotics. PLOS Biol. 2022 Aug 9;20(8):e3001721.

11. Li G, Zhang J, Guo Q, Jiang Y, Wei J, Zhao L li, et al. Efflux Pump Gene Expression in Multidrug-Resistant Mycobacterium tuberculosis Clinical Isolates. PLOS ONE. 2015 Feb 19;10(2):e0119013.

12. Henderson PJF, Maher C, Elbourne LDH, Eijkelkamp BA, Paulsen IT, Hassan KA. Physiological Functions of Bacterial “Multidrug” Efflux Pumps. Chem Rev. 2021 May 12;121(9):5417–78.

13. Jeannot K, Sobel ML, El Garch F, Poole K, Plésiat P. Induction of the MexXY Efflux Pump in Pseudomonas aeruginosa Is Dependent on Drug-Ribosome Interaction. J Bacteriol. 2005 Aug;187(15):5341–6.

14. Kang HW, Woo GJ. Increase of multidrug efflux pump expression in fluoroquinolone-resistant *Salmonella* mutants induced by ciprofloxacin selective pressure. Res Vet Sci. 2014 Oct 1;97(2):182–6.

15. Li XZ, Plésiat P, Nikaido H. The challenge of efflux-mediated antibiotic resistance in Gram-negative bacteria. Clin Microbiol Rev. 2015 Apr;28(2):337–418.

16. Schmalstieg AM, Srivastava S, Belkaya S, Deshpande D, Meek C, Leff R, et al. The Antibiotic Resistance Arrow of Time: Efflux Pump Induction Is a General First Step in the Evolution of Mycobacterial Drug Resistance. Antimicrob Agents Chemother. 2012 Aug 17;56(9):4806–15.

17. Vianna JS, Machado D, Ramis IB, Silva FP, Bierhals DV, Abril MA, et al. The Contribution of Efflux Pumps in Mycobacterium abscessus Complex Resistance to Clarithromycin. Antibiotics. 2019 Sept;8(3):153.

18. Rai D, Mehra S. The Mycobacterial Efflux Pump EfpA Can Induce High Drug Tolerance to Many Antituberculosis Drugs, Including Moxifloxacin, in Mycobacterium smegmatis. Antimicrob Agents Chemother. 2021 Oct 18;65(11):10.1128/aac.00262-21.

19. Silva KPT, Sundar G, Khare A. Efflux pump gene amplifications bypass necessity of multiple target mutations for resistance against dual-targeting antibiotic. Nat Commun. 2023 June 9;14(1):3402.

20. Rodríguez-Verdugo A, Gaut BS, Tenaillon O. Evolution of Escherichia coli rifampicin resistance in an antibiotic-free environment during thermal stress. BMC Evol Biol. 2013 Feb 22;13(1):50.

21. Huseby DL, Pietsch F, Brandis G, Garoff L, Tegehall A, Hughes D. Mutation Supply and Relative Fitness Shape the Genotypes of Ciprofloxacin-Resistant Escherichia coli. Mol Biol Evol. 2017 May 1;34(5):1029–39.

22. MacLean RC, Hall AR, Perron GG, Buckling A. The population genetics of antibiotic resistance: integrating molecular mechanisms and treatment contexts. Nat Rev Genet. 2010 June;11(6):405–14.

23. Yu XH, Hao ZH, Liu PL, Liu MM, Zhao LL, Zhao X. Increased Expression of Efflux Pump norA Drives the Rapid Evolutionary Trajectory from Tolerance to Resistance against Ciprofloxacin in Staphylococcus aureus. Antimicrob Agents Chemother. 2022 Nov 29;0(0):e00594–22.

24. Akanksha, Mehra S. Conserved Evolutionary Trajectory Can Be Perturbed to Prevent Resistance Evolution under Norfloxacin Pressure by Forcing *Mycobacterium smegmatis* on Alternate Evolutionary Paths. ACS Infect Dis. 2024 Aug 9;10(8):2623–36.

25. Chevereau G, Dravecká M, Batur T, Guvenek A, Ayhan DH, Toprak E, et al. Quantifying the Determinants of Evolutionary Dynamics Leading to Drug Resistance. PLOS Biol. 2015 Nov 18;13(11):e1002299.

26. Yu XQ, Yang H, Feng HZ, Hou J, Tian JQ, Niu SM, et al. Targeting efflux pumps prevents the multi-step evolution of high-level resistance to fluoroquinolone in Pseudomonas aeruginosa. Microbiol Spectr. 2025 Feb 21;13(4):e02981–24.

27. Papkou A, Hedge J, Kapel N, Young B, MacLean RC. Efflux pump activity potentiates the evolution of antibiotic resistance across S. aureus isolates. Nat Commun. 2020 Aug 7;11(1):3970.

28. Toprak E, Veres A, Michel JB, Chait R, Hartl DL, Kishony R. Evolutionary paths to antibiotic resistance under dynamically sustained drug selection. Nat Genet. 2012 Jan;44(1):101–5.

29. Costa SS, Viveiros M, Amaral L, Couto I. Multidrug Efflux Pumps in Staphylococcus aureus: an Update. Open Microbiol J. 2013 Mar 22;7:59–71.

30. Viveiros M, Martins A, Paixão L, Rodrigues L, Martins M, Couto I, et al. Demonstration of intrinsic efflux activity of *Escherichia coli* K-12 AG100 by an automated ethidium bromide method. Int J Antimicrob Agents. 2008 May 1;31(5):458–62.

31. Sparks IL, Derbyshire KM, Jacobs WR, Morita YS. Mycobacterium smegmatis: The Vanguard of Mycobacterial Research. J Bacteriol. 205(1):e00337–22.

32. Rodrigues L, Ramos J, Couto I, Amaral L, Viveiros M. Ethidium bromide transport across Mycobacterium smegmatiscell-wall: correlation with antibiotic resistance. BMC Microbiol 2011 111. 2011 Feb;11(1):1–10.

33. Pule CM, Sampson SL, Warren RM, Black PA, van Helden PD, Victor TC, et al. Efflux pump inhibitors: targeting mycobacterial efflux systems to enhance TB therapy. J Antimicrob Chemother. 2016 Jan 1;71(1):17–26.

34. Buroni S, Manina G, Guglierame P, Pasca MR, Riccardi G, De Rossi E. LfrR Is a Repressor That Regulates Expression of the Efflux Pump LfrA in Mycobacterium smegmatis. Antimicrob Agents Chemother. 2006 Dec;50(12):4044–52.

35. Elbourne LDH, Tetu SG, Hassan KA, Paulsen IT. TransportDB 2.0: a database for exploring membrane transporters in sequenced genomes from all domains of life. Nucleic Acids Res. 2017 Jan 4;45(D1):D320–4.

36. Beggs GA, Brennan RG, Arshad M. MarR family proteins are important regulators of clinically relevant antibiotic resistance. Protein Sci. 2020;29(3):647–53.

37. Birukou I, Tonthat NK, Seo SM, Schindler BD, Kaatz GW, Brennan RG. The Molecular Mechanisms of Allosteric Mutations Impairing MepR Repressor Function in Multidrug-Resistant Strains of Staphylococcus aureus. mBio. 2013 Aug 27;4(5):10.1128/mbio.00528-13.

38. Aguilar-Rodea P, Zúñiga G, Cerritos R, Rodríguez-Espino BA, Gomez-Ramirez U, Nolasco-Romero CG, et al. Nucleotide substitutions in the mexR, nalC and nalD regulator genes of the MexAB-OprM efflux pump are maintained in Pseudomonas aeruginosa genetic lineages. PLOS ONE. 2022 May 10;17(5):e0266742.

39. Schneiders T, Amyes SGB, Levy SB. Role of AcrR and RamA in Fluoroquinolone Resistance in Clinical Klebsiella pneumoniae Isolates from Singapore. Antimicrob Agents Chemother. 2003 Sept;47(9):2831–7.

40. Andries K, Villellas C, Coeck N, Thys K, Gevers T, Vranckx L, et al. Acquired Resistance of Mycobacterium tuberculosis to Bedaquiline. PLOS ONE. 2014 July 10;9(7):e102135.

41. Yang L, Shi H, Zhang L, Lin X, Wei Y, Jiang H, et al. Emergence of Two AcrB Substitutions Conferring Multidrug Resistance to Salmonella spp. Antimicrob Agents Chemother [Internet]. 2021 Mar 8 [cited 2025 Oct 1]; Available from: https://journals.asm.org/doi/10.1128/aac.01589-20

42. Zwama M, Nishino K. Ever-Adapting RND Efflux Pumps in Gram-Negative Multidrug-Resistant Pathogens: A Race against Time. Antibiotics. 2021 July;10(7):774.

43. Farzaneh S, Norouzi F, Fazeli H, Salehi M, Safari M, Esfahani BN. Novel mutation in efflux pump Rv1258c (Tap) gene in drug resistant clinical isolates of Mycobacterium tuberculosis in Iran. J Infect Dev Ctries. 2024 Feb 29;18(02):243–50.

44. Ma KC, Mortimer TD, Grad YH. Efflux Pump Antibiotic Binding Site Mutations Are Associated with Azithromycin Nonsusceptibility in Clinical Neisseria gonorrhoeae Isolates. mBio [Internet]. 2020 Aug 25 [cited 2025 Aug 25]; Available from: https://journals.asm.org/doi/10.1128/mbio.01509-20

45. Hawkey J, Ascher DB, Judd LM, Wick RR, Kostoulias X, Cleland H, et al. Evolution of carbapenem resistance in Acinetobacter baumannii during a prolonged infection. Microb Genomics. 2018;4(3):e000165.

46. Koutsolioutsou A, Peña-Llopis S, Demple B. Constitutive soxR mutations contribute to multiple-antibiotic resistance in clinical Escherichia coli isolates. Antimicrob Agents Chemother. 2005 July;49(7):2746–52.

47. Koutsolioutsou A, Martins EA, White DG, Levy SB, Demple B. A soxRS-Constitutive Mutation Contributing to Antibiotic Resistance in a Clinical Isolate of Salmonella enterica (Serovar Typhimurium). Antimicrob Agents Chemother. 2001 Jan;45(1):38–43.

48. Rodrigues L, Villellas C, Bailo R, Viveiros M, Aínsa JA. Role of the Mmr Efflux Pump in Drug Resistance in Mycobacterium tuberculosis. Antimicrob Agents Chemother. 2013 Jan 22;57(2):751–7.

49. Sander P, De Rossi E, Böddinghaus B, Cantoni R, Branzoni M, Böttger EC, et al. Contribution of the multidrug efflux pump LfrA to innate mycobacterial drug resistance. FEMS Microbiol Lett. 2000 Dec 1;193(1):19–23.

50. Rodrigues L, Ramos J, Couto I, Amaral L, Viveiros M. Ethidium bromide transport across Mycobacterium smegmatiscell-wall: correlation with antibiotic resistance. BMC Microbiol. 2011 Feb 18;11(1):35.

51. Lopatkin AJ, Bening SC, Manson AL, Stokes JM, Kohanski MA, Badran AH, et al. Clinically relevant mutations in core metabolic genes confer antibiotic resistance. Science. 2021 Feb 19;371(6531):eaba0862.

52. Puckett S, Trujillo C, Wang Z, Eoh H, Ioerger TR, Krieger I, et al. Glyoxylate detoxification is an essential function of malate synthase required for carbon assimilation in Mycobacterium tuberculosis. Proc Natl Acad Sci. 2017 Mar 14;114(11):E2225–32.

53. Serafini A, Tan L, Horswell S, Howell S, Greenwood DJ, Hunt DM, et al. Mycobacterium tuberculosis requires glyoxylate shunt and reverse methylcitrate cycle for lactate and pyruvate metabolism. Mol Microbiol. 2019 Oct;112(4):1284–307.

54. Bhat SA, Iqbal IK, Kumar A. Imaging the NADH:NAD+ Homeostasis for Understanding the Metabolic Response of Mycobacterium to Physiologically Relevant Stresses. Front Cell Infect Microbiol [Internet]. 2016 Nov 8 [cited 2025 Dec 8];6. Available from: https://www.frontiersin.org/journals/cellular-and-infection-microbiology/articles/10.3389/fcimb.2016.00145/full

55. Arce-Rodríguez A, Pankratz D, Preusse M, Nikel PI, Häussler S. Dual Effect: High NADH Levels Contribute to Efflux-Mediated Antibiotic Resistance but Drive Lethality Mediated by Reactive Oxygen Species. mBio. 2022 Jan 18;13(1):e02434–21.

56. Whittle EE, Orababa O, Osgerby A, Siasat P, Element SJ, Blair JMA, et al. Efflux pumps mediate changes to fundamental bacterial physiology via membrane potential. mBio. 2024 Sept 9;0(0):e02370–24.

57. Taneja NK, Tyagi JS. Resazurin reduction assays for screening of anti-tubercular compounds against dormant and actively growing Mycobacterium tuberculosis, Mycobacterium bovis BCG and Mycobacterium smegmatis. J Antimicrob Chemother. 2007 Aug;60(2):288–93.

58. Jadaun GPS, Agarwal C, Sharma H, Ahmed Z, Upadhyay P, Faujdar J, et al. Determination of ethambutol MICs for Mycobacterium tuberculosis and Mycobacterium avium isolates by resazurin microtitre assay. J Antimicrob Chemother. 2007 July 1;60(1):152–5.

59. Paixão L, Rodrigues L, Couto I, Martins M, Fernandes P, de Carvalho CCCR, et al. Fluorometric determination of ethidium bromide efflux kinetics in Escherichia coli. J Biol Eng. 2009 Oct;3(1):1–13.

60. Wright MH, Adelskov J, Greene AC. Bacterial DNA Extraction Using Individual Enzymes and Phenol/Chloroform Separation. J Microbiol Biol Educ. 2017 Sept;18(2):10.1128/jmbe.v18i2.1348.

61. Andrews, S. (2010) FastQC A Quality Control Tool for High Throughput Sequence Data. - References - Scientific Research Publishing [Internet]. [cited 2025 Nov 16]. Available from: https://www.scirp.org/reference/referencespapers?referenceid=2781642

62. Langmead B, Salzberg SL. Fast gapped-read alignment with Bowtie 2. Nat Methods. 2012 Apr;9(4):357–9.

63. Danecek P, Bonfield JK, Liddle J, Marshall J, Ohan V, Pollard MO, et al. Twelve years of SAMtools and BCFtools. GigaScience. 2021 Feb 1;10(2):giab008.

64. Koboldt DC, Zhang Q, Larson DE, Shen D, McLellan MD, Lin L, et al. VarScan 2: somatic mutation and copy number alteration discovery in cancer by exome sequencing. Genome Res. 2012 Mar;22(3):568–76.

65. Hunter JD. Matplotlib: A 2D Graphics Environment. Comput Sci Eng. 2007 May;9(3):90–5.

66. Harris CR, Millman KJ, van der Walt SJ, Gommers R, Virtanen P, Cournapeau D, et al. Array programming with NumPy. Nature. 2020 Sept;585(7825):357–62.

67. Waskom ML. seaborn: statistical data visualization. J Open Source Softw. 2021 Apr 6;6(60):3021.

68. McKinney W. Data Structures for Statistical Computing in Python. SciPy 2010 [Internet]. 2010 May 1 [cited 2025 Nov 16]; Available from: https://proceedings.scipy.org/articles/Majora-92bf1922-00a

69. The Galaxy Community. The Galaxy platform for accessible, reproducible, and collaborative data analyses: 2024 update. Nucleic Acids Res. 2024 July 5;52(W1):W83–94.

70. Liao Y, Smyth GK, Shi W. featureCounts: an efficient general purpose program for assigning sequence reads to genomic features. Bioinformatics. 2014 Apr 1;30(7):923–30.

71. Moderated estimation of fold change and dispersion for RNA-seq data with DESeq2 | Genome Biology | Full Text [Internet]. [cited 2025 Nov 16]. Available from: https://genomebiology.biomedcentral.com/articles/10.1186/s13059-014-0550-8

72. Kanehisa M, Furumichi M, Sato Y, Ishiguro-Watanabe M, Tanabe M. KEGG: integrating viruses and cellular organisms. Nucleic Acids Res. 2021 Jan 8;49(D1):D545–51.

73. Yu G, Wang LG, Han Y, He QY. clusterProfiler: an R package for comparing biological themes among gene clusters. Omics J Integr Biol. 2012 May;16(5):284–7.

74. R Core Team (2024) R A Language and Environment for Statistical Computing. R Foundation for Statistical Computing. - References - Scientific Research Publishing [Internet]. [cited 2025 Nov 16]. Available from: https://www.scirp.org/reference/referencespapers?referenceid=3887298

75. Singh A, Crossman DK, Mai D, Guidry L, Voskuil MI, Renfrow MB, et al. Mycobacterium tuberculosis WhiB3 Maintains Redox Homeostasis by Regulating Virulence Lipid Anabolism to Modulate Macrophage Response. PLOS Pathog. 2009 Aug 14;5(8):e1000545.

76. Vilchèze C, Weisbrod TR, Chen B, Kremer L, Hazbón MH, Wang F, et al. Altered NADH/NAD+ Ratio Mediates Coresistance to Isoniazid and Ethionamide in Mycobacteria. Antimicrob Agents Chemother. 2005 Feb;49(2):708–20.

77. Chawla M, Singh A. Detection of Membrane Potential in Mycobacterium tuberculosis. BIO-Protoc [Internet]. 2013 [cited 2025 June 17];3(11). Available from: https://bio-protocol.org/e785

